# Alternative splicing coupled nonsense-mediated decay shapes the temperature-dependent transcriptome

**DOI:** 10.1101/2020.02.19.956037

**Authors:** Alexander Neumann, Stefan Meinke, Gesine Goldammer, Miriam Strauch, Daniel Schubert, Bernd Timmermann, Florian Heyd, Marco Preußner

## Abstract

Mammalian body temperature oscillates with the time of the day and is altered in diverse pathological conditions. We recently identified a body temperature-sensitive thermometer-like kinase, which alters SR protein phosphorylation and thereby globally controls alternative splicing (AS). AS can generate mRNA variants containing premature termination codons, which are degraded by nonsense-mediated decay (NMD). Here we show extensive coupling of body temperature-controlled AS to NMD, leading to global control of temperature-dependent gene expression (GE). Temperature-controlled NMD-inducing splicing events are evolutionarily conserved and pervasively found within RNA-binding proteins, including most SR proteins. NMD-inducing exons are essential for rhythmic GE of SR proteins and have a global role in establishing temperature-dependent rhythmic GE profiles, both, in mammals under circadian body temperature cycles and in plants in response to ambient temperature changes. Together, these data identify body temperature-driven AS-NMD as an evolutionary ancient, core clock-independent mechanism to generate rhythmic GE.

## Introduction

Circadian clocks act as cell autonomous time-measuring devices, which anticipate active and resting phases and coordinate behavior and physiology accordingly. The mammalian circadian timing system is organized in a hierarchical manner: The central pacemaker in the brain’s suprachiasmatic nucleus (SCN) serves as a master regulator to coordinate circadian clocks throughout the body (Dibner, Schibler et al., 2010). Therefore, information on the geophysical time (light-dark) is transferred from the retina to the SCN, which then uses diverse neuronal and humoral cues to synchronize circadian clocks in other ‘light-blind’ parts of the brain and in other organs of the body (Gerber, Saini et al., 2015). Although the mechanisms of synchronization are different in the various organs, the conventional perception is that the majority of 24-hour rhythms depends on an identical transcription-translation feedback loop in each cell of the body (Ko & Takahashi, 2006). However, only ~50% of circadian mRNA rhythms depend on *de novo* transcription (Koike, Yoo et al., 2012, Menet, Rodriguez et al., 2012), strongly arguing for an involvement of other posttranscriptional mechanisms in generating rhythms (Preussner & Heyd, 2016, Shakhmantsir & Sehgal, 2019). Additionally, there is evidence for 24-hour rhythms which cycle independent of the central oscillator (Kornmann, Schaad et al., 2007). Core clock-independent 24-hour rhythms in GE may be difficult to discriminate from real circadian oscillations *in vivo* (reviewed in (Preussner & Heyd, 2018)) and our mechanistic understanding of these rhythms is limited.

Systemic body temperature cycles (Refinetti & Menaker, 1992) can synchronize the central oscillator in peripheral clocks (Brown, Zumbrunn et al., 2002, Buhr, Yoo et al., 2010, Saini, Morf et al., 2012), but can also directly result in rhythmic changes in GE. Examples include rhythmic expression of heat shock factors (Liu, Qian et al., 2019), of cold-induced RNA-binding proteins (RBPs), such as *Cirbp* (Liu, Hu et al., 2013, Morf, Rey et al., 2012), or rhythmic changes in the ratio of AS isoforms of *U2af26* and many other related genes (Goldammer, Neumann et al., 2018, Preussner, Goldammer et al., 2017, Preussner, Wilhelmi et al., 2014). All of these examples must be driven by highly sensitive mechanisms that are able to respond to 1-2°C changes in temperature (Gotic, Omidi et al., 2016, Preussner et al., 2017), but the global impact of these mechanisms (and more generally of body temperature) on GE patterns remains enigmatic so far. In our previous work, we revealed key components regulating temperature-dependent AS. Essential regulators are members of the family of SR proteins (13 canonical members in human), which share a domain rich in serine and arginine residues, known as the RS domain (Manley & Krainer, 2010). The phosphorylation level of SR proteins controls their activity and serves as a fast and extremely sensitive molecular thermometer transferring the signal ‘body temperature’ into global changes in AS (Haltenhof, Kotte et al., 2020). SR proteins contain ultraconserved elements, allowing autoregulation of their own expression level (Lareau, Inada et al., 2007). Autoregulatory mechanisms can involve regulated inclusion of an exon containing a premature translation termination codon (PTC; (Sureau, Gattoni et al., 2001)), or AS such that the normal stop codon becomes a PTC (Goncalves & Jordan, 2015, Lareau et al., 2007). PTC-containing mRNAs are recognized and degraded by the NMD surveillance pathway and for SR proteins this is understood as a mechanism associated with homeostatic control (Ni, Grate et al., 2007). However, the extreme evolutionary conservation is a strong indicator for further biological functions. AS-inducing NMD (AS-NMD) is not restricted to SR proteins and can thus dynamically control global GE (Braunschweig, Gueroussov et al., 2013, Lykke-Andersen & Jensen, 2015). While several thousand genes are affected by inhibition of the NMD pathway (Hurt, Robertson et al., 2013), the role of NMD in dynamically and actively controlling GE in response to changing cellular conditions has been described only in isolated cases (Tabrez, Sharma et al., 2017, Wong, Ritchie et al., 2013).

Here, we investigated to which extend AS-NMD is regulated by temperature and how this mechanism globally regulates body temperature-dependent GE. We find that NMD isoforms of different RBPs, especially SR proteins, are highly responsive to changes in the physiological temperature range, and that temperature-controlled AS-NMD of SR proteins is sufficient to *de novo* generate 24-hour rhythms in GE. Temperature-dependent AS-NMD is conserved from plants to human, controls thousands of mRNAs and represents a global mechanism for the generation of rhythmic 24-hour GE *in vivo*. The temperature-sensing feedback loop inducing AS-NMD bears striking similarity to the classical circadian transcription-translation feedback loop, both relying on rhythmic expression of a core machinery that controls itself and many output genes, but represents an independent and widespread mechanism to generate rhythmic GE. Furthermore, this mechanism shows wider evolutionary conservation than the classical core clock, pointing to an essential role in integrating temperature signals beyond the classical day-night cycle into cellular GE programs.

## Results

### Temperature-dependent AS-NMD occurs frequently in primary mouse hepatocytes

The detection of temperature-dependent, PTC-containing isoforms requires depletion or pharmacological inhibition of the NMD pathway (Rehwinkel, Raes et al., 2006). NMD occurs co-translational, and therefore blocking translation (e.g. via cycloheximide (CHX)) represents a fast and reliable way of blocking the NMD pathway (Hurt et al., 2013). To identify temperature-controlled splicing isoforms inducing NMD, we performed RNA-Seq on freshly-isolated primary mouse hepatocytes, incubated either at 34 or 38°C and treated either with DMSO or CHX (Figure 1A). A principal component analysis revealed clear clustering of the biological triplicates and comparable effects of both temperature and CHX on AS (Figure S1A). Using Whippet (Sterne-Weiler, Weatheritt et al., 2018), we initially identified over 3800 genes with CHX-sensitive splicing events at either 34°C or 38°C. A quarter of these events lie within genes previously associated with NMD (Figure S1B), validating our approach for identification of NMD-inducing isoforms. Next, we identified 4740 temperature-controlled (comparing 34°C and 38°C) splicing events in the CHX-treated cells (Figure S1C and Table S1). Only 1/3 of the strongest 1000 events and about 60% of all events respond in a similar manner in the DMSO control, indicating non-NMD events (Figures 1B and S1D, heat-skipped or cold-skipped). All other isoforms, almost 2000 in total (Table S1), lose their temperature-dependence in the DMSO control, indicating NMD-mediated degradation of one isoform, as it only accumulates in CHX-treated cells. These cases can be categorized into four groups: (I) cold-induced NMD via inclusion; (II) heat-induced NMD via inclusion; (III) cold-induced NMD via exclusion; and (IV) heat-induced NMD via exclusion. Examples for each of these categories were confirmed by radioactive RT-PCRs with RNAs from independently generated hepatocytes (Figures 1C and S1E) as follows: In *Mettl16*, exon 6 encodes a PTC, and the full length (fl) isoform – containing exon 6 – is therefore stabilized via CHX. Exon 6 inclusion is strongly promoted at 34°C, representing a cold-induced NMD event. In *Hnrnpdl*, exon 8 inclusion generates a second exon junction complex after the canonical stop in exon 7, turning this stop into a PTC (Lindeboom, Supek et al., 2016). The fl isoform is stabilized by CHX and, as it predominates at 38°C, represents a heat-induced NMD event. In *Hnrnph1* and *Slc9a8*, skipping of an exon (length not divisible by 3) generates a frameshift and a PTC in a downstream exon.

**Figure 1.**
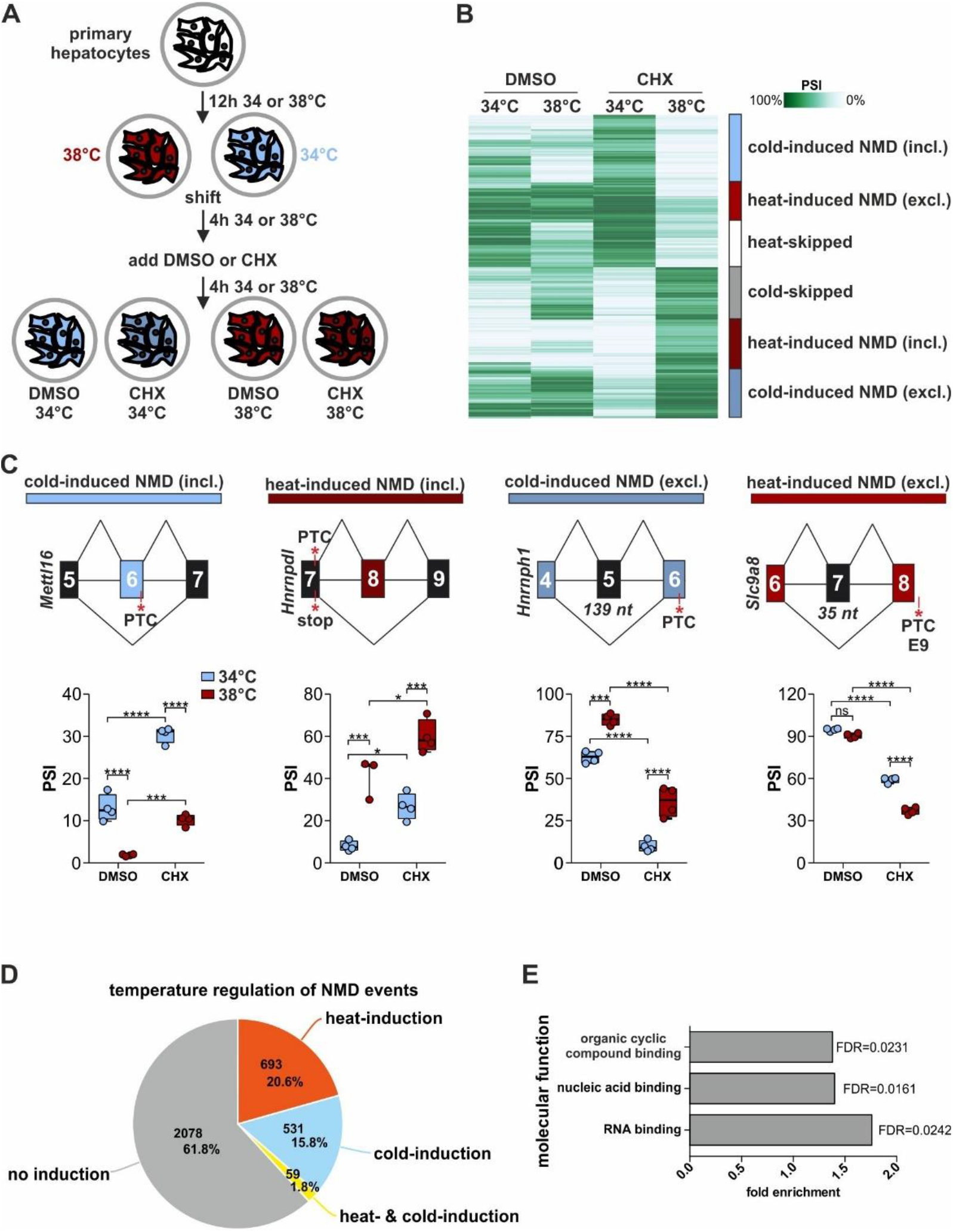
Characterization of temperature-controlled AS-NMD in mammalian cells. (A) Generation of RNA samples from mouse primary hepatocytes for analysis of temperature-dependent AS-NMD. (B) Automatic cluster-mapping of the temperature-dependent splicing changes in CHX, based on changes in PSI (percent spliced in). Categories (right) were manually assigned. Here the top 1000 events are shown, see also Figure S1B for all events. (C) Examples for cold-induced (blue) or heat-induced (red) AS-NMD via exon inclusion or exon skipping. In each example, on top a simplified exon-intron structure is given, PTCs and the length of frameshift inducing exons are highlighted. Predicted splicing changes were validated by radioactive RT-PCR with primers binding to the surrounding constitutive exons. Box plots show median PSI of validation PCRs of four different samples from at least two different mice, whiskers show min to max. Individual data points are shown as circles. Statistical significance was determined by 2-way ANOVA and is indicated by asterisks: adjusted p values: not significant (ns), *p<0.05, ***p<0.001, ****p<0.0001. See Figures S1C and S1E for further examples. (D) Classification of temperature-dependence in genes with NMD events. Number of genes and percentages are indicated. (E) GO-term analysis of temperature-controlled AS-NMD genes. All expressed genes (mean tpm > 0) served as background. See also Figure S1.

Globally, the NMD-inducing isoforms are dramatically stabilized by CHX, often with changes in NMD exon abundance of 60% or more, at either cold or heat conditions. In addition to skipped exon events, we also find temperature-dependent AS-NMD in the form of alternative 3’ or 5’ splice sites, intron retention, mutually exclusive exons, transcription starts and transcription ends (Table S1 and below), and the quantity and diversity of temperature-and CHX-controlled isoforms is indicative for a global impact on temperature-dependent GE levels. Additionally, certain NMD isoforms are almost exclusively found at one temperature (examples in *Rsrp1* and *Fus*; Figure S1E) and our approach of screening for AS-NMD at different temperatures thus expands the collection of known NMD-inducing isoforms.

Overall, inclusion levels (percent spliced in, PSI) predicted by Whippet and validated by RT-PCR from independent biological samples show strong correlation, allowing us to draw reliable conclusions from our bioinformatics analysis (Figure S1F). In all presented examples, as well as globally, the direction of PSI changes in DMSO correlate with PSI changes in CHX, confirming that these variants are present also in DMSO but are stabilized after CHX treatment (Figure S1G). When analyzing all genes with potential NMD events (all isoforms stabilized by CHX at 34 or 38°C) we find that almost 40% of these genes contain temperature-regulated NMD isoforms (Figures 1D and S1H), showing that NMD isoforms are pervasively responsive to temperature changes, which is consistent with a global role in shaping temperature-controlled GE. Interestingly, we find temperature-controlled NMD events enriched in RBPs (Figure 1E and below), which could represent the temperature-controlled core machinery that is able to amplify the temperature signal to control further downstream output events. In summary, temperature-controlled AS-NMD represents a frequent mode of posttranscriptional regulation, with wide implications for temperature-controlled GE levels.

### RBP expression is controlled through temperature-dependent AS-NMD

Our bioinformatics analysis identified over 60 RBPs with temperature-regulated NMD exons of different types (Figure 2A, see also Table S1). Most of these AS-NMD events are alternative cassette exons (e.g. *Hnrnpdl, Hnrnph3*), but there are also different types such as alternative last exons (e.g. *Cirbp*). In the presented examples, a temperature change of 4°C is sufficient to change AS by 20-30%, e.g. *Hnrnpdl* 60% to 95%, indicating a strong regulatory impact on the expression of the respective RBPs (Figure 2A). We have recently shown this particular temperature-dependent AS event in *Cirbp* to be regulated by CLK activity and found that E7b inclusion reduces *Cirbp* GE (Haltenhof et al., 2020). In line with these findings, we find the E7b isoform to be strongly stabilized by CHX and observe a correlation of higher abundance of the NMD isoforms (particularly in the CHX condition) with decreased overall GE in the DMSO control in *Cirbp* and all other cases analyzed (Figure 2B). For example, in *Hnrnpdl* the NMD isoform makes more than 90% at 38°C, correlating with a 4-fold reduction in GE. Note, that although the change in *Cirbp* exon 7b inclusion between the two CHX conditions is visible as a drastic change in the Sashimi plot (Figure 2A, only 10% of reads from exon 6 span to exon 7b at 34°C, while this is the case for almost 70% of reads at 38°C), Whippet quantifies this as a still severe but lower difference of about 15% (Figure 2B). For global correlations, we therefore include only skipped exon events, which are more robust to quantification. When considering all candidate RBPs with temperature-dependent AS-NMD skipped exon events, we observe a highly significant global correlation of temperature-dependent NMD-isoform generation and reduced GE levels in DMSO control (Figure 2C, left). Consistent with CHX preventing degradation of NMD-inducing isoforms, this correlation is completely abolished in the CHX samples (Figure 2C, right), strongly arguing for AS-NMD being the primary cause of temperature-dependent changes in GE levels. Looking at all genes with temperature-controlled cassette exon NMD events, the global trend is still present, although to a lesser extent (Figure S2), indicating that temperature-dependent GE requires AS-NMD but is additionally controlled by other mechanisms. In summary, these data reveal a mechanism that globally controls GE in a temperature-regulated manner via AS coupled to the NMD pathway.

**Figure 2.**
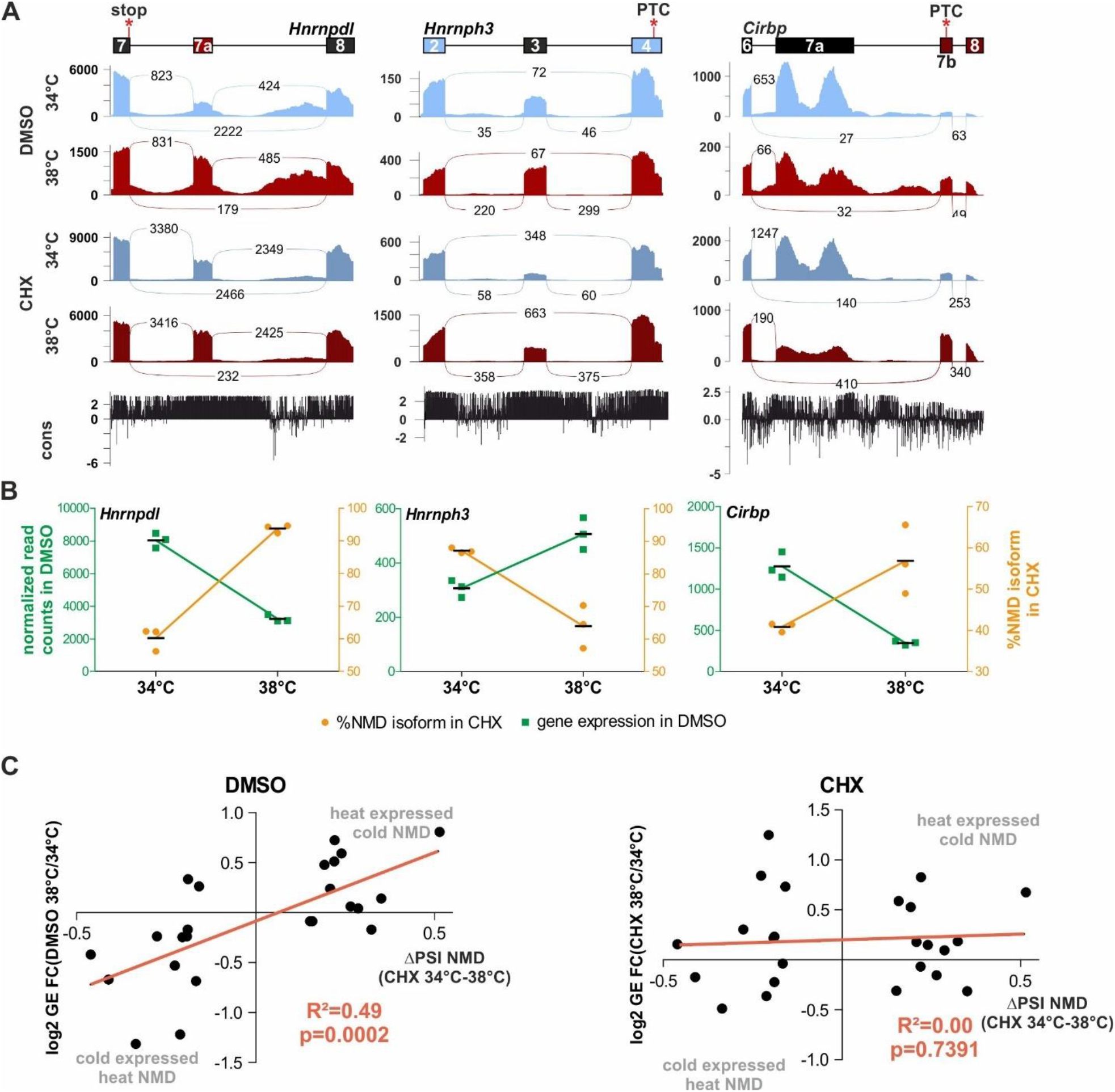
Temperature-dependent AS-NMD events globally regulate GE. (A) Temperature-dependent AS-NMD events in *Hnrnpdl* (left, see also Figure 1C), *Hnrnph3* (middle) and *Cirbp* (right). For each target, on top a simplified exon-intron structure is given and below Sashimi plots show the distribution of raw sequencing reads. Exon-Exon junction reads are indicated by the numbers connecting the exons. Below, sequence conservation across placental species is indicated. *Hnrnpdl* exhibits a heat-included exon that leads to a PTC, *Hnrnph3* a cold-skipped exon that leads to a frameshift, and in *Cirbp*, heat leads to inclusion of an alternative transcript end, which leads to formation of a PTC. (B) For each gene from A, the triplicate normalized read counts in DMSO (left y-axis, green) and the percentage of the NMD isoform in CHX (right y-axis, orange) are plotted at the two temperatures. (C) Correlation of NMD isoform inclusion and GE levels for all RBPs with temperature-dependent AS-NMD skipped exon events. Shown are the log2 fold change (FC) in GE versus the ΔPSI of the NMD isoform between the CHX samples. Left shows the GE change between the DMSO samples and right between the CHX samples. R^2^ and P (deviation from zero slope) are indicated. N=24. See also Figure S2.

### Temperature-controlled AS-NMD is evolutionarily conserved across endotherms

We next elucidated the evolutionary conservation of temperature-regulated AS-NMD. For SR proteins, it was previously described that these have NMD-inducing exons with ultraconserved regions (Lareau et al., 2007) but their function was only linked to producing homeostatic GE levels. We hypothesized that these ultraconserved regions likely have additional functionalities and could be involved in controlling temperature-dependent GE of SR proteins via AS-NMD. We indeed detect AS-NMD variants for most SR proteins, ranging from temperature-insensitive to strongly temperature-dependent (Figure S3A). Two extreme examples for heat-induced and cold-induced non-productive splicing events are ultraconserved exons in *Srsf2* and *Srsf10*. For *Srsf2* we observed and validated strong heat-induced inclusion of the PTC-encoding exon 3 (ΔPSI in CHX =46%, Figures 3A, left and S3B). In contrast, we observed almost cold-restricted generation of a non-productive isoform for *Srsf10*, namely exon 3 inclusion coupled to the use of an alternative splice site in exon 2 (ΔPSI in CHX =19%, Figures 3A, right and S3C), indicating antagonistic effects of temperature on different SR proteins. Again, the heat-induced NMD exon inclusion of *Srsf2* correlates with decreased GE in heat and vice versa for *Srsf10* (Figures 3A, center). To investigate whether temperature cycles could lead to rhythmic splicing of NMD inducing exons, we applied a modified simulated body temperature regime (Brown et al., 2002, Preussner et al., 2017) to human Hek293 cells. Consistent with our data from mouse hepatocytes, the corresponding AS events in *Srsf2* and *Srsf10* respond to 24-hour square-wave 34/38°C temperature rhythms in human Hek293 (Figure 3B), showing a cell-type-independent and evolutionarily-conserved regulation of SR proteins through (circadian) temperature changes. We also note that splicing-regulation is reverted already within 4-8 hours at one temperature, which is indicative for an autoregulatory feedback loop. Furthermore, we have validated additional temperature-dependent NMD isoforms for *Hnrnpdl*, *Hnrnph3*, and *Cirbp* in Hek293 cells (Figure 3C), consistent with high evolutionary conservation (see conservation track in Figure 2A). Importantly, we find that all temperature-controlled AS-NMD exons as well as several 100 nucleotides in the surrounding introns are extremely conserved across placental organisms (Figure 3D and conservation tracks in Sashimi plots in Figures 2A and 3A), arguing for an evolutionarily vital function. Consistent with (Thomas, Polaski et al., 2020), this conservation is present in all AS-NMD exons. Finally, we exemplarily investigated temperature-controlled *Srsf2* AS in hamster, rabbit and chicken cell lines (chicken as a non-placental endothermic organism), and strikingly find exon 3 inclusion promoted by heat and CHX in all cases (Figure 3E), thus demonstrating conserved regulation in all investigated endothermic organisms. The percentage of heat-induced NMD exon inclusion varies between 8% in rabbit cell or human Hek293 and up to 80% in primary mouse hepatocytes indicating species-or tissue-specific differences in SRSF10 expression or activity. In summary, we find that temperature-dependent NMD exon inclusion in SR proteins anticorrelates with SR protein expression and is highly evolutionarily conserved, indicating an essential function in regulating temperature-dependent GE.

**Figure 3.**
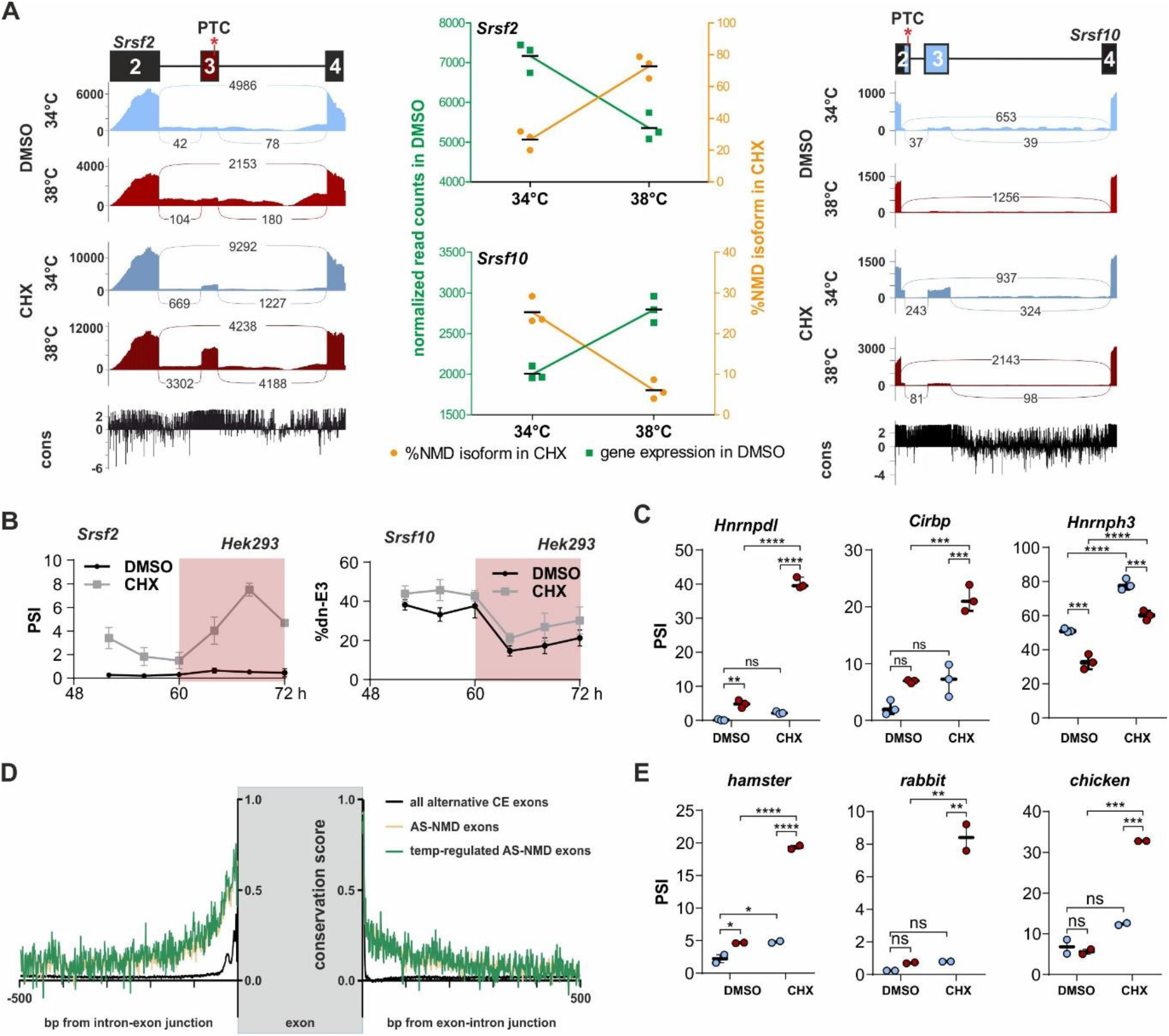
Temperature-dependent AS-NMD is evolutionarily conserved. (A) SR protein specific temperature-dependent AS-NMD. Sashimi plots are shown for heat-induced NMD exon inclusion for *Srsf2* (left) and cold-induced NMD isoform formation for *Srsf10* (right). In the center, quantification of GE and AS (as in Figure 2B). (B) NMD exon inclusion for *Srsf2* (left) and *Srsf10* (right) in a 24h temperature-rhythm in human cells. Hek293 cells were pre-entrained with square-wave temperature cycles (12h 34°C/ 12h 38°C) for 48h. For the last 24h cells were treated with DMSO or CHX every 4h and harvested after 4h and analyzed by splicing sensitive RT-PCR (n=3, mean ± SD). White area: 34°C; Red area: 38°C. In Hek293 cells, inclusion of *Srsf10* exon 3 is coupled to polyadenylation making it a weak NMD target. (C) AS-NMD of human targets at 34°C (blue) and 38°C (red) was investigated after a square-wave temperature regime (see B). RNAs were harvested after 56 and 68 hours, respectively. Statistical significance was determined by 2-way ANOVA and is indicated by asterisks: adjusted p values: not significant (ns), **p<0.01, ***p<0.001, ****p<0.0001. (D) Intron conservation of alternative cassette exons. Shown are average placental conservation scores introns surrounding all alternative exons (black, N=46,901), AS-NMD exons (yellow, N=453) and temperature-regulated AS-NMD exons (green, N=139). (E) Quantification of *Srsf2* AS-NMD in hamster, rabbit and chicken cells (n=2, mean ± SD). Statistical significance was determined by 2-way ANOVA, asterisks as in D, *p<0.05. See also Figure S3.

### Temperature rhythms are sufficient to generate GE rhythms depending on AS-NMD

To investigate whether external 24-hour rhythms in temperature are sufficient to generate rhythmic GE, we analyzed *Srsf2* and *Srsf10* GE in response to square-wave temperature rhythms (Figure 4A). Both genes clearly show 24-hour rhythms in gene expression, with *Srsf2* showing higher expression during the cold-phase (heat-induced NMD) and *Srsf10* showing higher expression during the warm phase (cold-induced NMD; WT in Figures 4B and 4C). Next, we generated CRISPR/Cas9-edited cell lines lacking either *Srsf2* exon 3 or *Srsf10* exon 3 and confirmed removal of the respective exons on RNA level (Figures S4A and S4B). Applying square-wave temperature rhythms to these cell lines, we find that removing the temperature-dependent exon is sufficient to abolish rhythms driven by external temperature cycles in both cases (ΔE3 in Figures 4B and 4C), showing that conserved NMD-inducing alternative exons in SR proteins indeed are necessary for their temperature-dependent GE.

**Figure 4.**
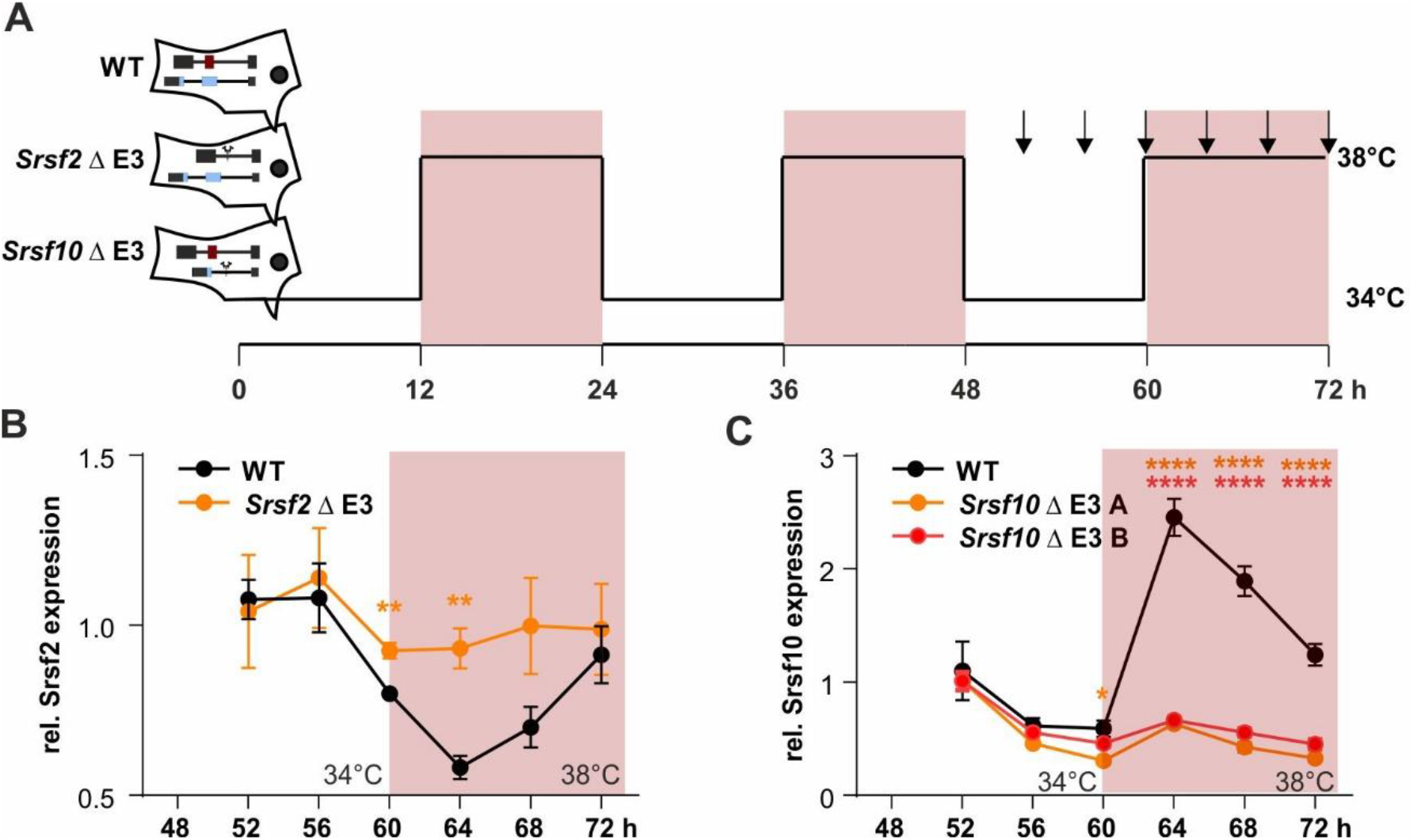
Temperature-dependent AS-NMD is necessary for rhythmic GE. (A) WT Hek293 and cell lines lacking either *Srsf2* or *Srsf10* exon 3 (ΔNMD Exon) through CRISPR/Cas9-mediated genome editing (see also Figure S4) where temperature entrained through 3 consecutive days of square-wave temperature rhythm (12 hours at 34°C and 38°C, respectively). Time points for harvest are indicated by arrows. (B, C) Rhythmic *Srsf2* (B) and *Srsf10* (C) productive mRNA levels (relative to *Gapdh*) were analyzed by RT-qPCR. For each clone, expression is normalized to time point 52h, (B) n = 3, (C) n =4, mean ± SEM. For *Srsf10* two independently generated clones were investigated (ΔE3 A and B). Statistical significance was determined by unpaired t-test and is indicated by asterisks: p values: *p<0.05, **p<0.01, ****p<0.0001. See also Figure S4.

### Temperature-dependent AS-NMD regulates expression levels of SR proteins in *A. thaliana*

The pervasive identification of temperature-controlled AS-NMD in distinct SR proteins (and other RBPs) indicates an evolutionary advantage in homeotherms. To span a broader evolutionary spectrum, we next asked whether a similar mechanism is also in place for poikilotherms, which experience much greater differences in (body) temperature (10°C during a day-night cycle and often 50°C in seasonal changes compared to 1-2°C change in body temperature in mice). We therefore examined temperature-dependent SR protein splicing and expression in *A. thaliana*, encoding 18 SR proteins (based on (Barta, Kalyna et al., 2010)) with evolutionary conserved regions triggering AS-NMD (Palusa & Reddy, 2010, Rauch, Patrick et al., 2014, Richardson, Rogers et al., 2011). We analyzed a publicly-available RNA-Seq time course dataset from plants kept at 20°C for 24 hours and then shifted to 4°C for the next 24 hours, while being under a 12-hour dark-light regime (Calixto, Guo et al., 2018), allowing us to distinguish between light-and temperature-dependent effects. When assessing changes in GE, we strikingly find that all 18 SR proteins show a clear temperature-dependent regulation (Figure 5A). Some genes additionally show a light-dependent expression pattern, but temperature is in general the dominant signal. About half of the regulated SR proteins show higher GE at 20°C, while the other half is cold-induced. Interestingly, we find at least one example for both warm-and cold-expressed SR genes in all but one SR protein subfamilies (Figure 5B), which may suggest partially complementing functions. Within the RNA-Seq data we find clear evidence for NMD-inducing AS that is antagonistically regulated to GE in 50% of temperature regulated SR proteins (Figures S5A and S5B), for example in At-SR34 and At-SCL33 (Figure 5C), consistent with previous notions (Calixto et al., 2018, Staiger & Brown, 2013). In all tested examples, temperature-dependent splicing and GE changes could be confirmed by RT-PCR from independently-generated plant samples (Figures 5D and S5C). To systematically characterize temperature-dependent AS-NMD events in plants, we first used RNA-Seq data of CHX-treated plants to identify NMD events (Drechsel, Kahles et al., 2013) and then quantified temperature regulation of these targets in samples without NMD inhibition (data from (Calixto et al., 2018)). Although detection of NMD isoforms is challenging without stabilization of the respective isoforms, we were able to identify 80 NMD isoforms clearly affected by temperature (Table S1). Consistent with our findings in mouse, there is a highly significant anti-correlation between NMD isoform inclusion and GE (Figure 5E). This includes a temperature-controlled AS-NMD event in the snRNP assembly factor At-GEMIN2 (Figure S5D), which was previously identified as a key regulator of downstream temperature-dependent clock-related splicing events (Schlaen, Mancini et al., 2015). In conclusion, these data expand our findings from the mammalian to the plant system, identifying temperature-dependent AS-NMD as an evolutionary pervasive mechanism that adapts SR protein expression to changing temperatures. During mammalian evolution, AS-NMD in SR genes arose independently, evolutionary rapid and in an SR protein specific manner (Lareau & Brenner, 2015), thus showing that *cis*-regulatory elements regulating temperature-dependent AS-NMD independently arose in homeotherms and poikilotherms. This suggests a broad evolutionary advantage of temperature-dependent SR protein expression and is indicative for an evolutionary-conserved and-adapted temperature sensor controlling SR protein activity.

**Figure 5.**
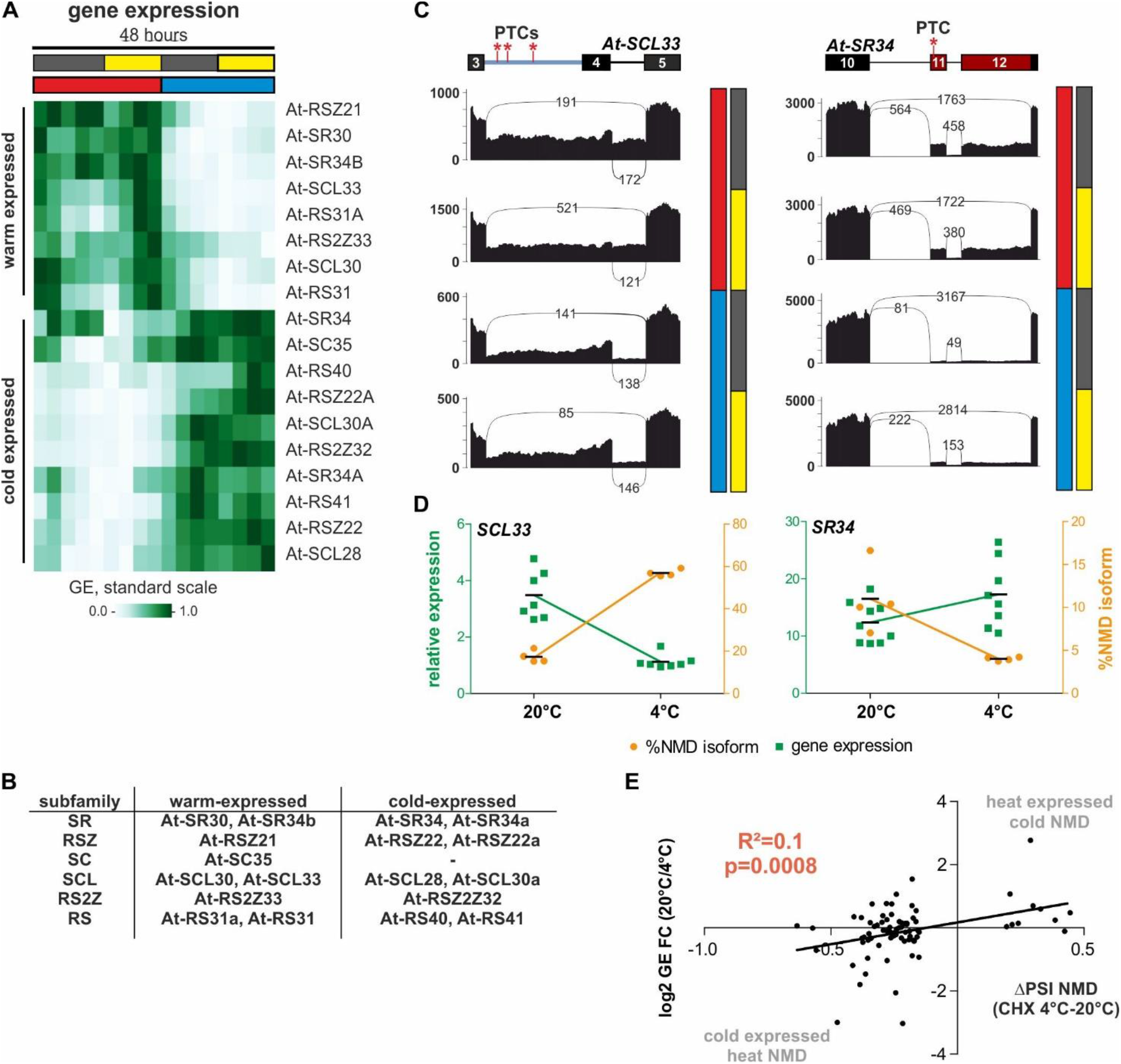
Temperature-dependent AS-NMD of SR proteins is conserved in plants. (A) Normalized GE values of plant SR proteins in a 48-hour time course. In the first day, plants were kept at 20°C (red) and shifted to 4°C (blue) for the second day. Plants were under a 12-hour dark/light regimen (gray/yellow bars). Warm-and cold-expressed genes are indicated. (B) *A. thaliana* SR proteins divided in their subfamilies and classified as warm-or cold-expressed. (C) Sashimi plots as shown in Figure 2A (without conservation score) for NMD-inducing AS events upon cold (At-SCL33, left) or warm temperature (At-SR34, right). Data shown represent, from top to bottom, warm-dark, warm-light, cold-dark and cold-light conditions as indicated on the right of the plots. (D) For validation *A. thaliana* were incubated for 3 days either at 20°C or 4°C and RNA was investigated for gene expression (green, left y-axis) and NMD isoform formation (orange, right y-axis). Gene expression of At-SCL33 (left) or At-SR34 (right) is shown relative to Ipp2. (E) Correlation of NMD isoform inclusion and GE levels for temperature-dependent AS-NMD skipped exon events. Shown are the log2 fold change (FC) in GE versus the ΔPSI of the NMD isoform between the CHX samples. R^2^ and P(deviation from zero slope) are indicated. N=80 (see main text for details). See also Figure S5.

### Body temperature-regulated AS-NMD generates 24-hour rhythms in GE

Finally, we address the question to which extent rhythmic GE *in vivo* is depending on naturally-occurring body temperature cycles. We therefore compared the acrophase (time of maximal GE) of heat and cold-induced genes across a circadian cycle in mouse liver (Atger, Gobet et al., 2015). Strikingly, we find that genes with higher expression at 38°C in hepatocytes predominantly peak during the night when mice are active and have a higher body temperature. In contrast, cold-induced genes from hepatocytes rather peak during the colder day (Figure 6A, top), providing evidence for a direct function of body temperature cycles in generating 24-hour rhythmic GE. Consistent with a function of temperature-dependent AS-NMD in this mechanism, we additionally find that genes with a cold-induced NMD skipped exon event peak during the night, while genes with a heat-induced NMD skipped exon event tend to peak during the day (Figure 6A, bottom). Of note, of the 254 genes with temperature-induced AS-NMD skipped exon events, 102 were found to exhibit cyclic expression.

**Figure 6.**
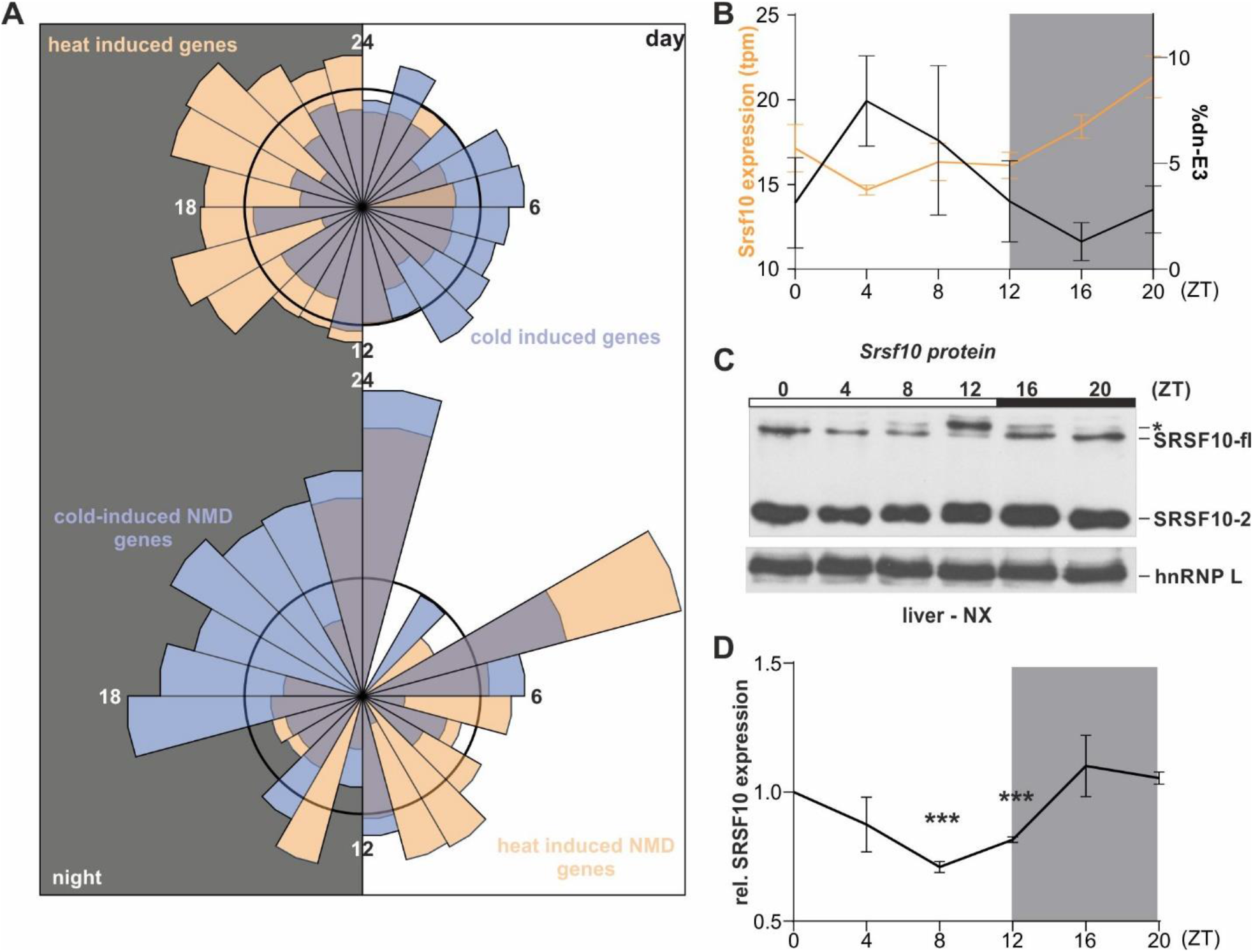
AS-NMD generates GE rhythms *in vivo*. (A) Top: Rose plot visualizing the acrophases (based on rhythmic genes in the liver transcriptome from (Atger et al., 2015)) of genes exhibiting heat-induced (orange, n=797) and cold-induced (blue, n=828) upregulation of GE in mouse hepatocytes, respectively. Bottom: Acrophases of genes with cold-induced (blue, n=55) and heat-induced (orange, n=47) skipped exon NMD events, respectively. Numbers indicate respective ZT, with the left half representing night (at which mice are active and have an elevated body temperature) and the right half representing day (at which mice are asleep with lower body temperature). The black circles represent expected normal distribution if data would be randomly sampled. (B) Correlation of rhythmic *Srsf10* expression (orange, left y-axis) and exon 3 inclusion (black, right y-axis) in mouse liver samples from the indicated ZTs (n=4, mean +/-SEM). Quantifications were obtained by RNA-seq analysis of datasets from (Atger et al., 2015). (C) Representative Western blot of SRSF10 protein levels from mouse liver nuclear extracts (NX) from different ZTs. The different SRSF10 variants are highlighted on the right. hnRNP L was used as a loading control. The asterisks could represent hyperphosphorylated SRSF10-fl through higher CLK activity during the day. (D) Quantification of SRSF10-fl + SRSF10-2 relative to ZT0 and hnRNP L (mean of at least 3 mice ± SEM). Student’s unpaired t test-derived p values ***p<0.001. See also Figure S6.

As one particular example, we analyzed *Srsf10* expression *in vivo* in detail and find a clear time of the day-dependent formation of the non-productive *Srsf10* isoform (Figure 6B, right y-axis). Consistent with our data from cultured hepatocytes, we observe antagonistic 24-hour rhythms in GE across a circadian day *in vivo* in liver (Figure 6B, left y-axis). This is not a tissue-specific but a general effect, as shown by validations in samples from mouse cerebellum confirming increased non-productive splicing in combination with lower GE during the day (Figure S6A). Similar to other temperature-driven AS events (Preussner et al., 2014), we find that *Srsf10* AS persists in constant darkness and is entrainable to a new light-dark regime (Figures S6B and S6C), highlighting common, e.g. body temperature-dependent, regulatory principles. Finally, we found that AS-NMD driven rhythms in GE are also reflected at the protein level by confirming daytime-dependent SRSF10 protein expression in mouse liver (Figures 6C and 6D). In summary, these data point to a fundamental role of mammalian body temperature in generating rhythmic GE *in vivo*, which is independent from the classical, circadian clock-mediated generation of 24-hour rhythms (Figure 7).

**Figure 7.**
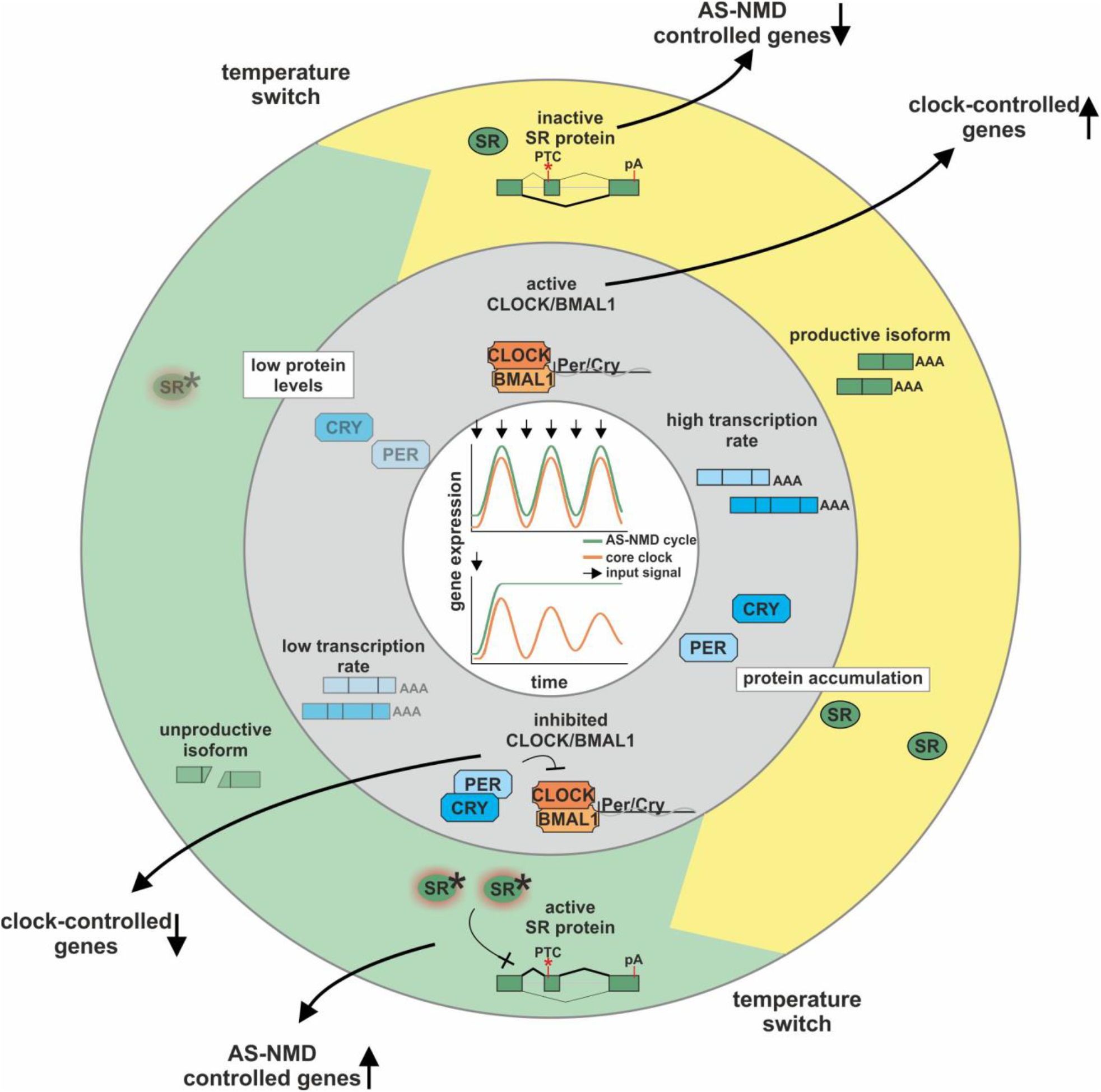
A model of two independent cycles generating rhythmic GE. Inner ring (grey): Classical transcription-translation feedback loop or the central circadian clock, involving the transcription activator CLOCK and BMAL1 and the transcription targets/repressors *Period* (Per) and *Cryptochrome* (Cry). Outer ring (green/yellow): Model of rhythmic GE driven via AS-NMD. AS regulates the formation of productive stable or unproductive instable isoforms, thereby regulating the abundance of SR protein mRNA and protein. An additional regulatory layer is achieved through temperature-mediated control of SR protein activity (active SR proteins are indicated by an asterisks), resulting in rhythmic GE of SR proteins themselves and output target genes. GE in response to continuous (top) or single (bottom) input signal (indicated by arrows) is shown in the center.

## Discussion

Here we report that temperature-regulated AS coupled to NMD occurs frequently in diverse genes from evolutionary distant species. We furthermore show that these splicing events are sufficient to generate temperature-dependent GE changes and to induce 24-hour rhythms in GE. Together, these data are consistent with a model where body temperature-controlled AS-NMD represents a second, core clock-independent generator of 24-hour rhythmic GE. Classical circadian cycles in GE are driven by a transcription-translation feedback loop (Figure 7, inner ring). Here, transcriptional activation trough CLOCK/BMAL1 results in accumulation of PERIOD (PER) and CRYPTOCHROME (CRY). When a critical protein level is reached, these factors dimerize, enter the nucleus and repress their own GE. This results in low mRNA and protein expression, again allowing activation through CLOCK and BMAL1. Together with *Per* and *Cry*, many other Clock-controlled genes oscillate through this feedback loop. Circadian oscillations are robust to environmental changes and continue to oscillate with only mildly-affected amplitude in the absence of rhythmic stimuli (Figure 7, center). In a second and largely independent cycle, body temperature induces rhythmic GE through AS-NMD (Figure 7, outer ring). Here, body temperature cycles control the activity and AS-NMD-mediated autoregulation of SR proteins. An SR protein that is inactive due to its phosphorylation level results in the accumulation of its mRNA and (inactive) protein. Upon a switch to the activating body temperature, phosphorylation of the SR protein changes through altered CLK activity (Haltenhof et al. 2020) and the activated SR proteins promotes NMD exon inclusion (within their own pre-mRNA). Degradation of the unproductive mRNA isoform then leads to low but active protein levels. Only upon a second switch to the inactivating temperature, predominant formation of the productive isoforms becomes possible, again resulting in accumulation of the respective SR protein. As for the classic circadian feedback loop, autoregulation of SR proteins represents the core machinery, which then controls other genes containing NMD exons to also cycle in response to rhythmic body temperature changes. As long as the body temperature oscillates, AS-NMD-driven GE also cycles in a circadian-like manner (Figure 7, center). This however requires continuous input and the rhythmicity is lost under constant conditions, which sets it apart from classic circadian GE.

In addition to SR proteins, we identified temperature-controlled NMD-inducing splicing isoforms in many other RBPs. Mechanistically, it will be interesting to investigate, whether these splicing events are directly controlled by SR proteins or whether temperature also controls the activity of other RBPs (e.g. by phosphorylation or other post-translational modifications). The presence of temperature-regulated AS-NMD isoforms in many RBPs is indicative for an interlocked network of RBPs to post-transcriptionally adapt GE to temperature, representing a potential layer of signal amplification. In mammals, this network requires sensors and modulators that are sensitive enough to respond to temperature differentials of around 1°C, to be able to respond to daily changes in body temperature. In recent work we have characterized CLK kinases as the direct molecular sensor of such temperature changes (Haltenhof et al., 2020). Interestingly we find in evolutionary distant species, that groups of SR proteins are antagonistically-affected by temperature, such that one group is activated by hyperphosphorylation, whereas this leads to inactivation of another group of SR proteins. While we observe this regulatory pattern in evolutionary distant species such as plants and humans, pointing to an important functionality, it remains unknown, how phosphorylation of SR proteins can have opposing effect on their activity. Autoregulatory feedback loops were initially characterized as a mechanism allowing homeostatic regulation of SR proteins (Lareau et al., 2007, Ni et al., 2007). Here, we report that the same autoregulatory feedback loop is capable of producing rhythms in GE. An input signal controlling

SR protein activity is sufficient to change SR protein expression. This has also intriguing consequences for SR protein target genes, as activation of a protein reduces total levels of the same protein. Target genes with high-affinity binding sites might therefore be bound and activated (including the own pre-mRNA) while target genes with low affinity binding sites are not bound, due to the reduced total protein levels, and therefore rather repressed.

At first sight the existence of two independent 24-hour rhythms appears puzzling, but these two rhythms are fundamentally different: While classical circadian rhythms depend on transcription-translation feedback loops and are robust to minor ambient changes (Gonze, Halloy et al., 2002), temperature-induced rhythms are generated post-transcriptionally and can quickly adapt GE in response to changing temperatures. Therefore, temperature-induced rhythms can stabilize circadian rhythms under circadian temperature cycles (Morf et al., 2012) but can additionally adapt GE in response to sudden temperature changes, i.e. under disease-associated stress conditions, and are therefore able to integrate all temperature-changing signals into complex GE programs. In mammals, this could example involve body temperature changes associated with the female hormone cycle (Nagashima, 2015) or with aging (Keil, Cummings et al., 2015). Furthermore, the interplay of these two mechanisms could be involved in seasonal control of GE by integrating light and temperature signals. For example, At-RS40 exhibits an intriguing upregulation of GE in cold light conditions (Figure 5A) and could therefore realize specific effects during winter days.

Homeothermic and poikilothermic organisms experience quite different differentials in body temperature, and it is therefore a striking finding that in mouse/human (1-2°C) and plants (>20°C) an almost identical mechanism controls SR protein levels. The evolution of AS-NMD in SR proteins (Lareau & Brenner, 2015) indicates an evolutionary independent origin of the temperature-controlled exons/introns. Mechanistically it will therefore be interesting to investigate how cis-regulatory elements, SR proteins and kinases are evolutionary adapted to allow control of SR protein expression in the body temperature range of diverse organisms. The evolutionary-pervasive identification of temperature-controlled AS-NMD in SR proteins is indicative for an important function in temperature adaptation. Additionally, the fact that ultraconserved elements within SR proteins have evolved rapidly and are effectively ‘frozen’ in mammals and birds (Bejerano, Pheasant et al., 2004), which are all endothermic homeotherms, represents a strong connection to endothermy. Because of the high sensitivity of temperature-controlled AS-NMD, this mechanism could be involved in setting or maintaining a constant body temperature. The clinical importance of ultraconserved exons triggering NMD is furthermore highlighted by their potential tumor suppressive function (Thomas et al., 2020) and the association of NMD isoforms with neuronal diseases (Jaffrey & Wilkinson, 2018). Therefore, the *de novo* identification of temperature-restricted NMD isoforms might reveal novel candidates for antisense oligonucleotide-based therapies in human (Bennett, Krainer et al., 2019). Overall, we describe a surprisingly widespread mechanism that integrates diverse environmental signals to induce AS-NMD cycles and generate GE rhythms from plants to human, constituting a plethora of future implications.

## Materials & Methods

### Mouse maintenance

All animal experiments were performed with C57BL/6 mice in accordance with institutional and governmental recommendations and laws. Mice were kept under constant 12-hour light-dark conditions. RNA samples across a circadian day, from constant darkness or after jet-lag were previously generated (Preussner et al., 2014). Mice of both genders were used. For preparation of RNA, tissues were quickly removed, frozen in liquid nitrogen and homogenized in RNATri (Bio&Sell). For preparation of nuclear extracts, we first prepared single cell suspensions of freshly isolated liver samples.

### Tissue culture cells

The HEK293T cell line has been present in the lab for over 5 years and is maintained in liquid nitrogen. Early passage aliquots are thawed periodically. Rabbit RK-13, Chinese hamster CHO and chicken enterocyte 8E11 cell lines were a kind gift from Dusan Kunec (FU Berlin). For all cell lines, cell morphology and growth is routinely assessed and corresponds to the expected phenotype. Cell cultures are tested for mycoplasma contamination monthly using a PCR-based assay. HEK293T and 8E11 cell lines were maintained in DMEM medium containing 10% FBS and Pen/Strep (Invitrogen). RK13 cells were maintained in EMEM medium containing 10% FBS and Pen/Strep (Invitrogen). CHO cells in Ham’s F12 medium containing 10% FBS and Pen/Strep. All cell lines were usually maintained at 37°C and 5% CO^2^, except for 8E11 cells (39°C). For square-wave temperature cycles we used two incubators set to 34°C and 38°C and shifted the cells every 12 hours. See (Preussner et al., 2017) for an image of the square-wave temperature cycles with temperatures and time points. For chicken cells we used 34°C and 40°C (as 37°C degrees would already be a reduced temperature). Transfections of Hek293T using Rotifect (Roth) were performed according to the manufacturer’s instructions. Cycloheximide (Sigma) was used at 40μg/ml final concentration or DMSO as solvent control.

To isolate primary mouse hepatocytes, liver was perfused with PBS and digested using Collagen digestion solution. Liver was transferred into a Petri dish and cells were liberated by mechanical force. Cells were washed three time with Williams Medium E and 10% FCS and plated.

### A. thaliana

To confirm temperature-dependent alternative splicing and gene expression in *A. thaliana* we isolated RNA from 28-day old Col-0 plants kept only at 20°C or 31-day old Col-0 plants kept at 4°C for the last 3 days. For RNA isolation using Trizol (see below) we used ~100mg material from leaves.

### RT-PCR and RT-qPCR

RT-PCRs were done as previously described (Preussner et al., 2014). Briefly, RNA was extracted using RNATri (Bio&Sell) and 1μg RNA was used in a gene specific RT-reaction. For analysis of minigene splicing the RNA was additionally digested with DNase I and re-purified. Low-cycle PCR with a ^32^P-labeled forward primer was performed, products were separated by denaturing PAGE and quantified using a Phosphoimager and ImageQuantTL software. For qRT-PCR up to 4 gene-specific primers were combined in one RT reaction. qPCR was then performed in a 96 well format using the ABsolute QPCR SYBR Green Mix (Thermo Fisher) on Stratagene Mx3000P instruments. qPCRs were performed in duplicates, mean values were used to normalize expression to a housekeeping gene (mouse: *Hprt*, human: *Gapdh*, plants: *Ipp2*; ΔCT) and Δ(ΔCT)s were calculated for different conditions. See Table S2 for primer sequences.

### RNA-Seq analysis and bioinformatics

Mapping of reads to reference genomes was performed using STAR version 2.5.3a (Dobin, Davis et al., 2012). Reference genomes mm10 (mouse) and TAIR10 (*A*. thaliana) were applied. Gene expression analysis was performed using Salmon version 0.11.3 transcript quantification (Patro, Duggal et al., 2017) followed by DESeq2 version 1.22.2 quantification (Love, Huber et al., 2014) both used according to their documentation.

Whippet version 0.11 (Sterne-Weiler et al., 2018) was used to obtain splicing ratios and transcript per million GE quantifications. To obtain splicing quantifications of AS-NMD events, index creation was supplemented with mapped reads of NMD-inhibited sequencing samples (see Table S1 for plant accession numbers) and the low TSL flag was not set. A splice event was considered significantly different between two conditions with a |ΔPSI| > 15%, probability > 85% and on average more than 10 junction reads. For AS-NMD events of SR proteins that could not be properly quantified by Whippet, PSI values were calculated based on junction read counts (e.g. Srsf7). Downstream analyses were performed using Python3, bash and R scripts. Most relevant Python packages used were pandas (general secondary data analysis, (McKinney, 2010)), numpy (numerical operations, (Walt, Colbert et al., 2011)), Matplotlib (data visualization, (Hunter, 2007)) and scikit-learn (principle component analysis, (Pedregosa, Varoquaux et al., 2011)). Molecular function analysis was performed using PANTHER version 14.1 (Mi, Muruganujan et al., 2013) with all genes expressed above 1 TpM as background. Sashimi plots were generated using a customized version of ggsashimi (Garrido-Martin, Palumbo et al., 2018), which additionally displays conservation scores. Junction reads with low count numbers were removed for clarity. Phylogenetic p-value conservation score data of placental organisms was downloaded from UCSC (mm10 phyloP60way placental, (Pollard, Hubisz et al., 2010, Siepel, Bejerano et al., 2005)) and manipulated with BEDOPS version 2.4.26 and bedtools version 2.26.0 (Neph, Kuehn et al., 2012, Quinlan & Hall, 2010).

For mouse data, splicing events were classified into the following groups:

1. events that show differential splicing upon temperature (34°C vs 38°C between DMSO or CHX conditions)
2. events that show differential splicing upon CHX treatment (DMSO vs CHX in 34°C or 38°C conditions) and no temperature-dependent differential splicing (34°C vs 38°C in CHX)
3. events that show differential splicing upon CHX treatment (DMSO vs CHX in 34°C or 38°C conditions) and additional temperature-dependent differential splicing (34°C vs 38°C in CHX)
4. events that do not fall into any of the above categories

Splicing events in groups 2) and 3) were identified as AS-NMD events. The temperature-regulated events were further divided into cold-induced and heat-induced AS-NMD events (considering directionality of the CHX-induction). AS-NMD events that showed different directionality upon CHX treatment at the two temperatures were omitted from the analysis (these made up <2% of CHX-induced events).

For *A. thaliana*, AS-NMD events were identified by using independent RNA-seq data from CHX-treated plants. These data did not include a temperature shift and therefore the classification between temperature-dependent and-independent AS-NMD events has been done using temperature shifted but not CHX treated data. Otherwise the classification was performed as for mouse.

For preparation of rose plots, gene acrophases (time of highest expression) from (Atger et al., 2015) were used and combined with information about temperature-regulation on expression (based on DESeq2 analysis, padj < 0.05) and AS-NMD level (based on Whippet and post-analysis, see above). Genes were sorted into 24 bins that represent 1h each. As the data are highly skewed, normalization was needed. Therefore, the percentage of genes falling into each bin was calculated for all genes obtained from (Atger et al., 2015), and the subsets of heat-expressed and cold-expressed genes as well as those of genes with cold-induced and heat-induced NMD events. The percentages for each bin in the subsets was then divided by this bin’s percentage of overall genes. The black circles shown in Figure 6A represent the normalization line (expected proportion of genes if no effect was present).

### Generation of CRISPR/Cas9 modified cell lines

For genome-engineering in Hek293T cells, sequences flanking the conserved exons of Srsf2 or Srsf10 (human) sgRNA candidates *in silico* using the Benchling tool. Upstream and downstream of each exon at least one pair of oligos for the highest ranked candidate sgRNA (Ran, Hsu et al., 2013) was synthesized and subcloned into the PX459 vector (kindly provided by Stefan Mundlos). sgRNA sequences are available on request. Cells were transfected in 6-well plates using Rotifect following the manufacturer’s protocol. 48 hours after transfection, the transfected cells were selected with 1 μg/ml puromycin and clonal cell lines were isolated by dilution (Ran et al., 2013). Genomic DNA was extracted using DNA extraction buffer (200 mM Tris pH 8.3, 500 mM KCl, 5 mM MgCl2, 0.1% gelatin in H2O) and a PCR was performed using gene-specific primers to confirm the exon knockout on DNA level. In promising clones the exon knockout was additionally confirmed after RNA isolation by splicing sensitive PCR.

### Western Blot

Nuclear fractionations (NX) were performed as previously described (Heyd & Lynch, 2010). SDS-PAGE and Western blotting followed standard procedures. Western blots were quantified using the ImageQuant TL software. The following antibodies were used for Western blotting: hnRNP L (4D11, Santa Cruz), SRSF10 (T-18, Santa Cruz).

### Quantification and statistical analysis

Figure legends contain information on repetitions and statistical tests used.

### Data and Software Availability

Full gel images and raw data files will be deposited as Mendeley Data. RNA sequencing data are available under GEO XXX.

### Contact for Reagent and Resource Sharing

Please contact M.P. for reagents and resources generated in this study. Accession numbers to all RNA-seq datasets used in this study are noted in Table S1.

## Acknowledgments

The authors would like to thank Jessica Stock, Alena Lohnert, Kevin Huolt and Hoonsung Cho for their contribution to this work as rotation students. Furthermore, we would like to thank Dirk Hincha for discussing AS-NMD in plants, Tom Haltenhof for the isolation of primary hepatocytes, the SeqCore facility for generation of RNA Sequencing data and the HPC Service of ZEDAT, Freie Universität Berlin, for computing time. This work was funded through DFG grants HE5398/4-2 and 278001972 - TRR 186 to FH. MP is funded by a stipend from the Peter Traudl Engelhorn Foundation.

## Author Contributions

SM, GG, MS and MP performed experiments. AN performed the bioinformatics analysis. DS provided plant material. BT performed RNA-sequencing. AN, MP and FH designed the study, planned experiments, analyzed data and wrote the manuscript with help from SM. MP and FH conceived and supervised the work.

## Supplemental information

**Figure S1.**
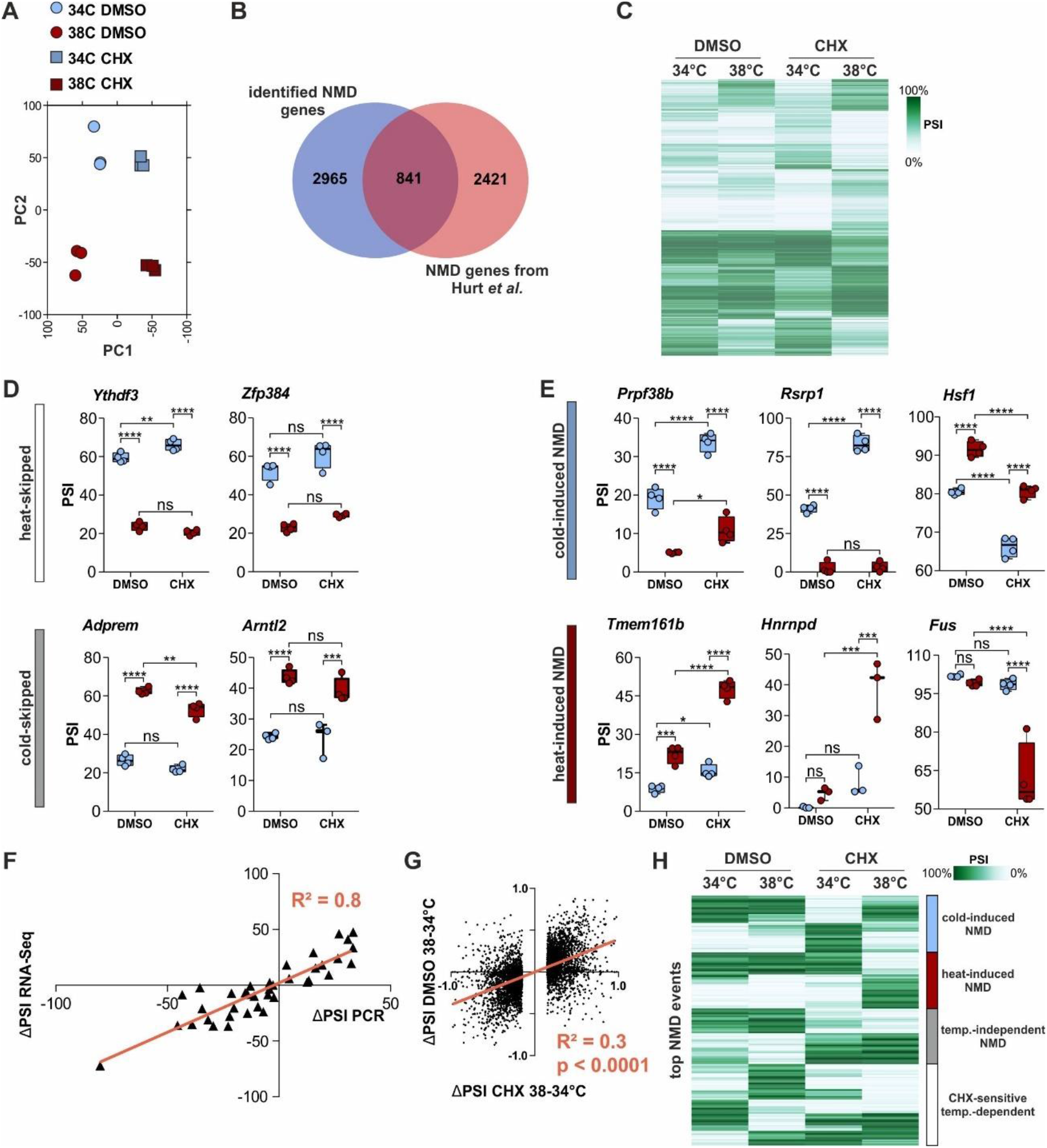
Validation of temperature-dependent AS in mammalian cells. (A) Principle component analysis of the investigated triplicate samples by RNA sequencing. (B) Overlap of genes with NMD events identified in our study in primary mouse hepatocytes (34 and 38°C) and by Hurt et al., 2013 in embryonic stem cells (37°C). (C) Automatic cluster-mapping of temperature-dependent splicing changes in CHX, based on PSI (percent spliced in). All events with |ΔPSI|>0.15 and p>0.85 between CHX 34 and 38°C are shown. N=4740. (D) Examples for heat-skipped (white) or cold-skipped (grey) splicing events. Analysis as in Figure 1C. Note that these splicing events are barely affected by CHX. (E) Further examples for cold-induced (blue) or heat-induced (red) AS-NMD events. Analysis as in Figure 1C. (F) Comparison of temperature-regulated AS events predicted by Whippet RNA-Seq analysis and validated by RT-PCR from independent samples. The line depicts linear regression, goodness of fit is represented by R^2^. (G) Comparison of splicing changes in CHX (x-axis) and DMSO (y-axis) for all events presented in S1 B. The line depicts linear regression, goodness of fit is represented by R^2^, N=4740. The p value represents statistical probability of the slope being different from 0. Splicing changes in DMSO and CHX have the same directionality but ΔPSI values are generally larger in CHX. (H) Temperature dependence of the strongest NMD events (top 500, largest ΔPSI in either 34 or 38°C comparison of DMSO vs CHX). Categories (right) were manually assigned. We assign four categories: (I) cold-induced NMD, (II) heat-induced NMD, (III) temperature independent NMD, and (IV) CHX-sensitive temperature-dependent splicing events. The generation of these isoforms could depend on *de novo* protein synthesis.

**Figure S2.**
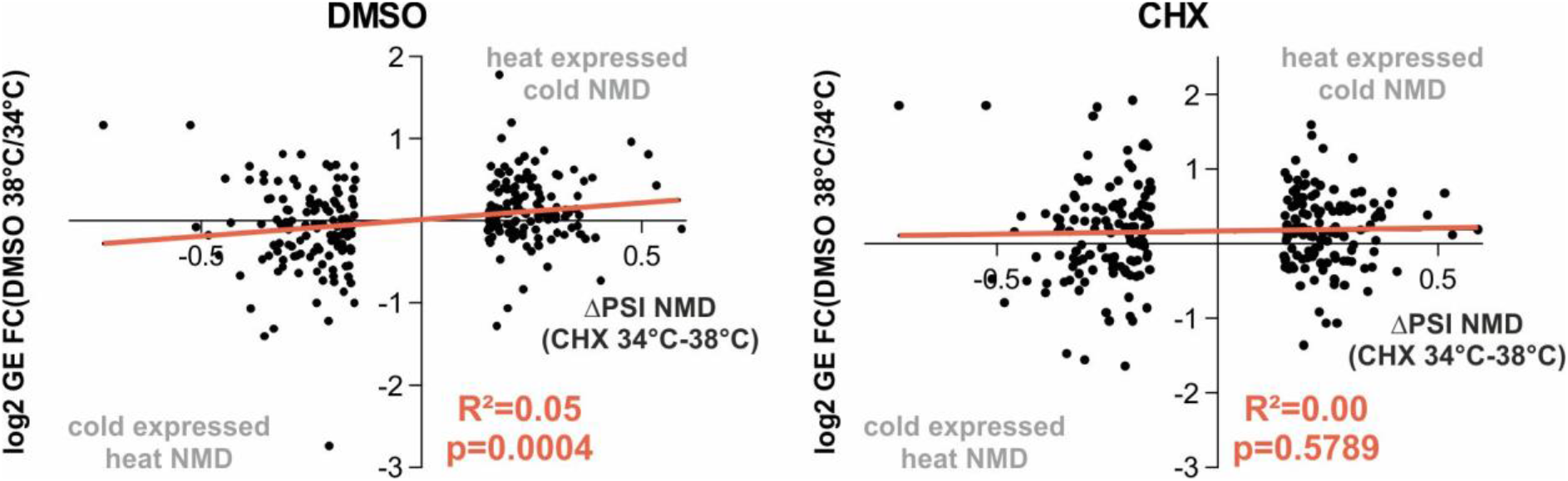
Temperature-dependent AS-NMD events globally regulate GE. Global correlation of NMD isoform inclusion and GE levels for all genes with temperature-dependent AS-NMD skipped exon events. Shown are the log2 fold change (FC) in GE versus the ΔPSI of the NMD isoform between the CHX samples. Left shows the GE change between the DMSO samples and right the between the CHX samples at the two temperatures. R^2^ and P (deviation from zero slope) are indicated, N=254.

**Figure S3.**
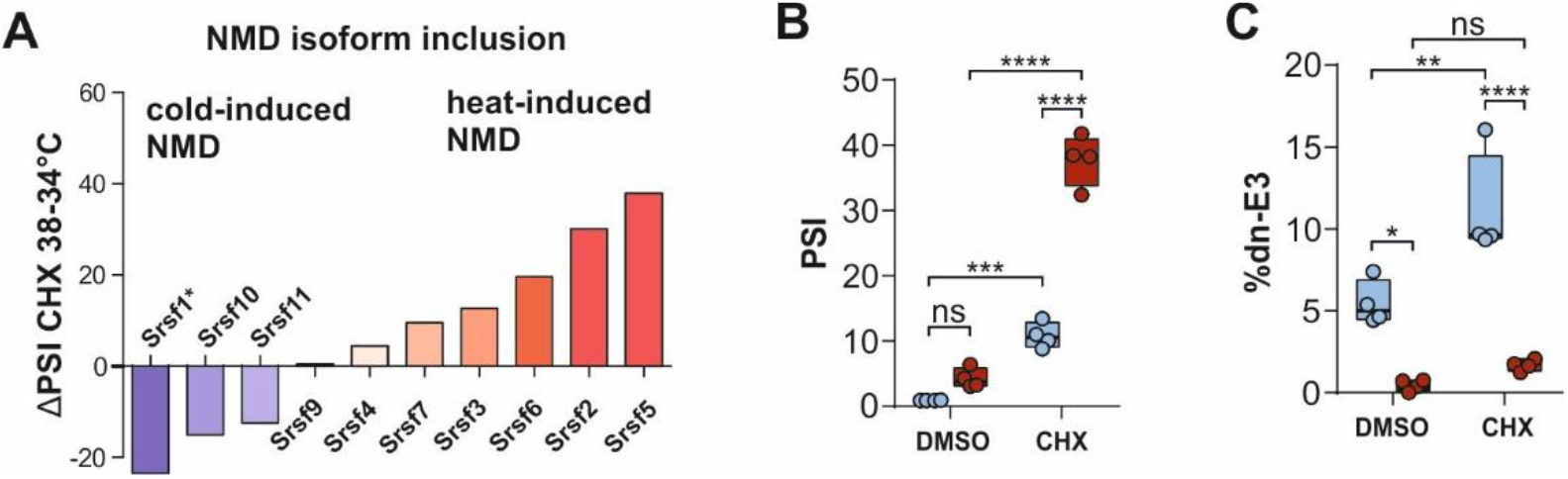
Temperature-dependent AS-NMD is evolutionary conserved. (A) Comparing temperature-dependent AS-NMD (ΔPSI of 38°C-34°C in CHX) of SR proteins in primary mouse hepatocytes. The NMD event in *Srsf1* (*) escaped the computational analysis; data are derived from radioactive RT-PCR. (B) Validation of *Srsf2* temperature-dependent AS-NMD by radioactive RT-PCR as described in Figure 1C. Statistical significance was determined by 2-way ANOVA and is indicated by asterisks: adjusted p values: not significant (ns), ***p<0.001, ****p<0.0001. (C) Validation of *Srsf10* temperature-dependent AS-NMD by radioactive RT-PCR as described in A. Asterisks as in B, *p<0.05.

**Figure S4.**
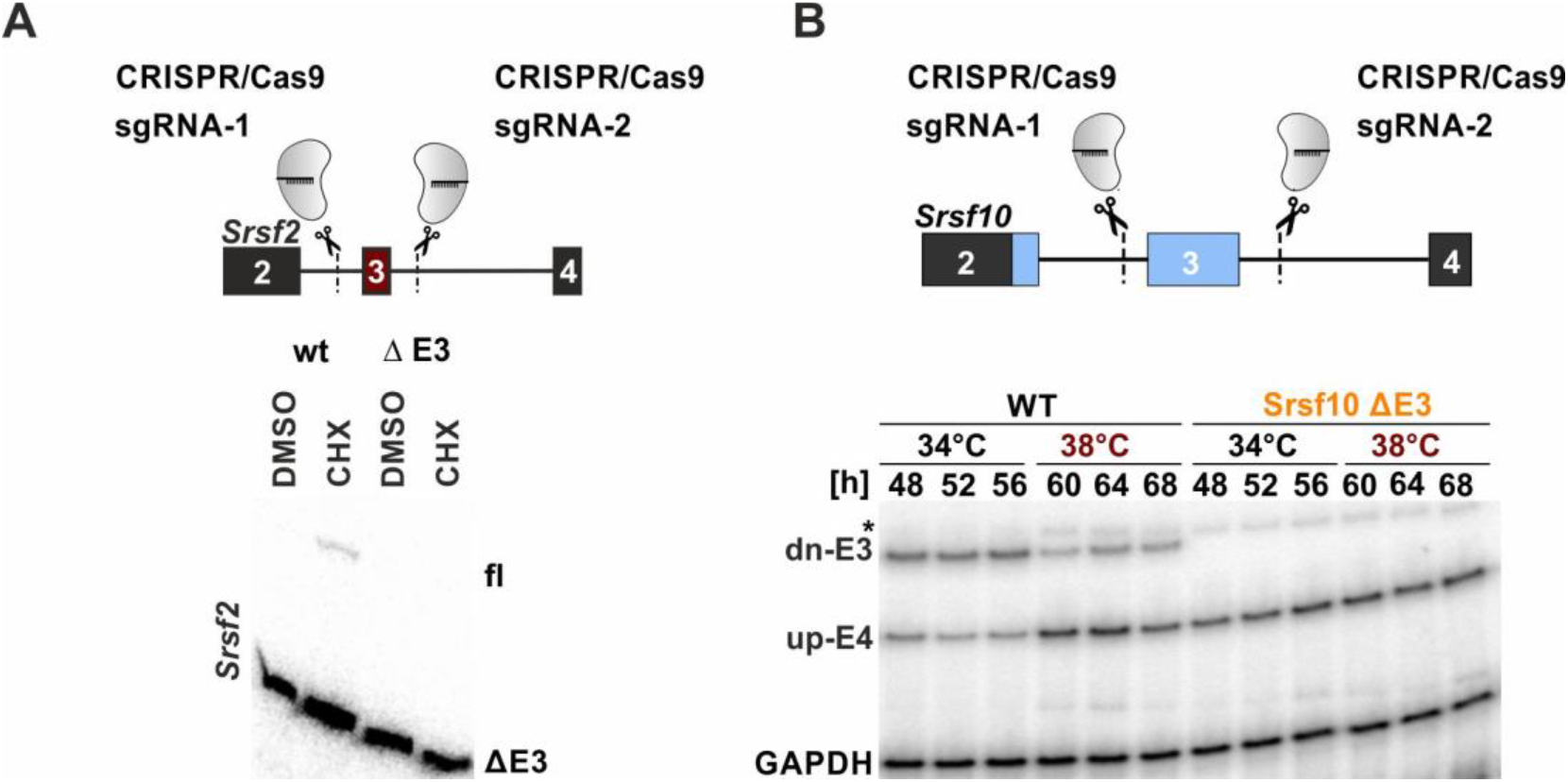
Temperature-dependent AS-NMD is necessary for rhythmic GE. (A, B) CRISPR/Cas9 strategy for cell lines lacking NMD exons for *Srsf2* (A) and *Srsf10* (B). Lack of NMD exons in clonal *Srsf2* CRISPR/Cas9-edited cell lines was confirmed by splicing sensitive RT-PCR (bottom). WT and mutant cell lines were treated with DMSO or CHX at 38°C and inclusion of the NMD isoform was investigated by splicing sensitive RT-PCR (A). Investigation of *Srsf10* dn-E3 splicing in samples from Figure 4C. AS was analyzed by RT-PCR. A *Gapdh* PCR was performed simultaneously and served as a loading control. A representative gel image is shown. The asterisk indicates an unspecific product.

**Figure S5.**
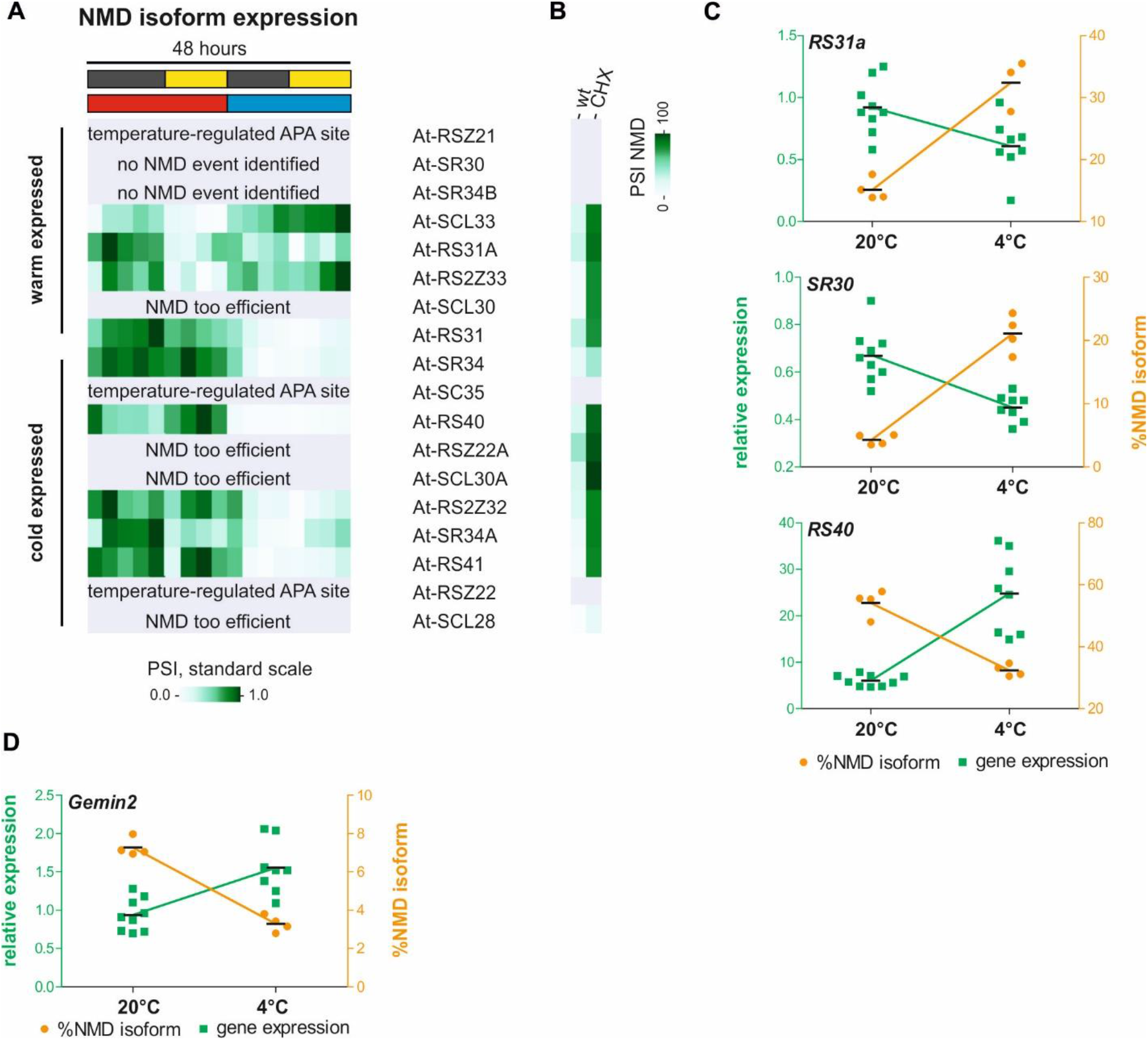
Temperature-dependent AS-NMD of SR proteins is conserved in plants. (A) Normalized NMD isoform expression of plant SR proteins as in Figure 5A. In this dataset, without inhibition of the NMD pathway, the identification of NMD isoforms is likely incomplete (compare to B). In three cases, a clear temperature-regulated alternative poly-adenylation (APA) event occurred but was not quantified (also indicating a post-transcriptional origin of temperature-dependent GE). In two cases, no NMD event could be identified. Four times NMD seemed to be too efficient, as no events could be identified without CHX treatment (B) AS-NMD isoform inclusion in wt or CHX-treated plants for the same genes as in A. Here the amount of the respective isoforms was compared in control and CHX treated samples (based on another RNA-Seq dataset (Drechsel et al., 2013), see Table S1). Note that most temperature depend isoforms from A are indeed stabilized upon CHX treatment. (C) Validation RT-PCRs and qPCRs of NMD events in further plant SR proteins as shown in Figure 5D. For each case, NMD isoform inclusion correlates with lower GE. (D) Validation of an AS-NMD event in Gemin2, in which cold-induced inclusion of the NMD isoform correlates with lower GE.

**Figure S6.**
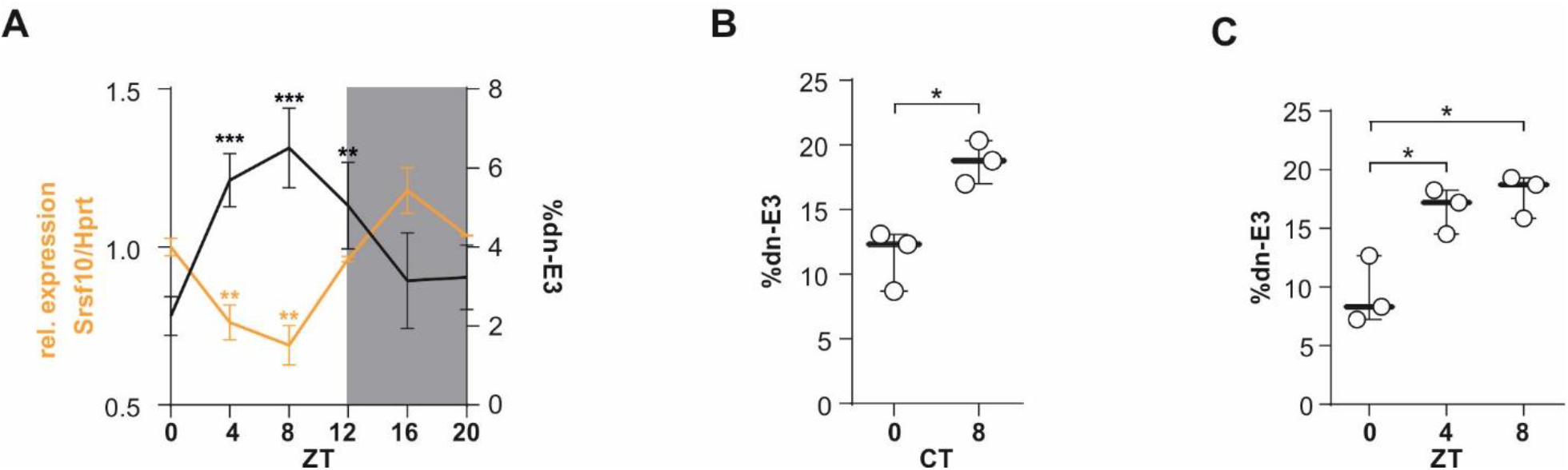
AS-NMD generates GE rhythms *in vivo*. (A) Correlation of rhythmic productive *Srsf10* mRNA expression (orange, left y-axis) and exon 3 inclusion (black, right y-axis) in mouse cerebellum samples from the indicated ZTs. Splicing was analyzed using radioactive RT-PCR of at least 3 mice per time point. Expression was determined by RT-qPCR and is shown relative to *Hprt* and ZT0 (mean of at least 3 mice ± SEM). Student’s unpaired t test-derived p values **p<0.01 indicate difference from ZT0. (B) Rhythmic *Srsf10* AS persists in constant darkness. Mice were kept in constant darkness for 24h and sacrificed at the indicated circadian times (CTs) of the following subjective day. RT-PCR analysis as in A (n=3, mean ± SD). Student’s unpaired t test-derived p values *p<0.05. (C) Entrained *Srsf10* AS. Mice were 8-hour phase delayed and on the 4th day sacrificed at the indicated ZTs. RT-PCR analysis as in A (n=3, mean ± SD). Student’s unpaired t test-derived p values *p<0.05.

